# Interpretable morphology mapping of peripheral blood leukocytes using annotation-efficient artificial intelligence

**DOI:** 10.64898/2026.05.22.725537

**Authors:** Zhuohe Liu, Simon P. Castillo, Xin Han, Xiaoping Sun, Zhihong Hu, Yinyin Yuan

## Abstract

**Background:** Peripheral blood smears (PBS) review is labor-intensive, subjective, and challenging for rare or morphologically heterogeneous cell types in hematologic malignancies. Artificial intelligence (AI) offers a scalable alternative, but broader clinical translation is constrained by annotation burden and limited interpretability.

**Methods:** We developed an interpretable, annotation-efficient AI framework that learns leukocyte morphology through a two-stage process: label-free representation learning to construct a morphological embedding space, followed by supervised fine-tuning for cell type and morphological attribute classification. The model was trained and evaluated on 5,952 PBS images from cancer patients at MD Anderson Cancer Center, including blast cells, and 17,092 images from public sources. Active learning strategies were assessed to improve label efficiency, and interpretability was examined using saliency and embedding visualization. An interactive web application, HemoSight, was developed to support clinical review.

**Findings:** The framework achieved a macro-F_1_ score of 0·96 for 9-way leukocyte classification on the internal test split and 0·83 on the held-out patient cohort. Active learning substantially reduced annotation requirements, reaching peak performance with only 13·3% of available labels and significantly improving learning efficiency across 8 of 9 cell types. The model generalized to classifying 11 leukocyte morphological attributes with a mean F_1_ score of 85·8% and revealed structured morphological landscapes. Saliency maps, embedding visualizations, and the HemoSight application enabled transparent morphological inspection of model predictions, supporting confidence in model behavior and feasibility for clinical integration.

**Interpretation:** Our framework enables scalable, annotation-efficient, and interpretable modeling of leukocyte morphology, supporting the integration of AI-assisted PBS review for hematopathology workflows.

**Funding:** Seed funding from The University of Texas MD Anderson Cancer Center.

**Research in Context:** *Evidence before this study:* Peripheral blood smear review is essential for diagnosing and monitoring hematologic malignancies, but manual case review is time-consuming and variable, particularly for rare or abnormal leukocyte types. Automated hematology analyzers are widely used to flag abnormal cells; however, they provide limited morphological insight and often require frequent manual correction, especially in cancer settings where disease and treatment alter cell appearance. Previous artificial intelligence approaches for leukocyte classification have shown promise, but most rely on fully supervised learning, require extensive expert annotation, focus on a limited set of cell types, and frequently exclude diagnostically important rare cells such as blasts. Interpretability is inconsistently addressed, and few studies provide tools that allow clinicians to inspect and interpret model outputs within routine workflows.

*Added value of this study:* This study introduces an annotation-efficient framework trained on a large collection of peripheral blood smear images, including cancer patient samples with hematopathologist-verified rare cell types such as blasts. The framework learns leukocyte morphology from unlabeled images and adapts to multiple classification tasks with minimal expert labeling. Performance is evaluated on both internal test splits and a held-out patient cohort to provide a realistic estimate of generalization. Iterative, uncertainty-guided annotation substantially reduces labeling requirements while improving learning efficiency across most leukocyte classes. Beyond cell-type classification, the framework is extended to 11 clinically relevant morphological attributes and reveals a structured morphological landscape. These capabilities are integrated into a web application, HemoSight, enabling real-time inference and transparent morphological inspection of predictions within hematopathology workflows.

*Implications of all the available evidence:* Advancing artificial intelligence for hematology requires methods that reduce expert labeling demands, provide interpretable outputs, and perform reliably across clinically diverse patient samples. This study shows that learning from largely unlabeled data combined with iterative expert annotation can support scalable and flexible modeling of leukocyte morphology for classification tasks. Integrating quantitative predictions and interactive visualization supports the use of artificial intelligence as an assistive tool for diagnostic peripheral blood smear review, with potential to improve efficiency, consistency, and reviewer confidence.

## Introduction

Peripheral blood smear (PBS) is a routine laboratory examination essential for screening and diagnosing of many hematological malignancies [1]. PBS enables the assessment of the morphological diversity among leukocytes, with their population and abundance serving as key indicators of disease stage [2,3]. Furthermore, leukocyte cytomorphology is important for the evaluation of treatment response and disease progression [4]. The wide adoption of automated analyzers has modernized hematology laboratory workflows, where PBS cases are routinely flagged by analyzers before manual review [5]. However, the slow turnaround time of manual PBS reviews creates bottlenecks, potentially delaying critical diagnoses of hematological disorders such as acute promyelocytic leukemia (APL), which can lead to life-threatening outcomes [6].

Despite their advantages, current hematology analyzers rely heavily on manual reviews and lack in-depth morphological analysis. These systems digitize regions of smear slides, typically capturing images at low magnifications to identify leukocytes before acquiring high-magnification images for cell classification [7]. This process is sensitive to clinical conditions such as clumping or hyperleukocytosis, leading to misclassification [8]. Misclassification is particularly pronounced in cancer hospitals, where treatment protocols and cancer-related conditions, including anemia and myelosuppression, necessitate frequent manual corrections [9]. Additionally, current analyzers have limited explainability, leaving many cytomorphological features unevaluated [10]. This limitation restricts the number of identified cell types and hinders usability. Because clinical significance depends on precisely defined morphological abnormalities, manual evaluations remain indispensable [11]. These drawbacks underscore the need for innovative approaches that enhance cell classification in PBS without compromising flexibility or interpretability.

Many efforts have been made to improve automated leukocyte classification, leveraging artificial intelligence (AI) to account for image heterogeneity [12,13]. In research settings, most approaches, relying on supervised learning, struggle to generalize to rare cell types and lack mechanisms to incorporate expert reclassification to improve future performance. Moreover, supervised learning demands extensive datasets with well-defined classes. Yet, many studies focus on a limited number of cell types, often excluding blasts or blast equivalents [14–16], including for example, monoblasts and promonocytes, which are crucial for diagnosing acute leukemia with monocytic differentiation [17]. To reduce false positives, some models incorporate pseudo-cell type classes, such as smudge cells or artifacts, which further increase annotation burden [18]. These limitations, combined with the regulatory and safety requirement for locked systems, translate to slow adoption and performance improvement of clinical AI applications. Concordance with expert annotation, especially on rare and abnormal cell types [19], have ample room of improvement despite decades of incorporating neural networks into commercial analyzers [7], and post-classification manual verification continues to enhance diagnostic accuracy [20,21].

Among AI-based methods, label-efficient and adaptive self-supervised learning (SSL) has emerged to transform automation in medical imaging and computational pathology [22,23]. SSL leverages unlabeled data, substantially reducing annotation costs [24]. When combined with active learning, SSL models can further improve label efficiency by progressively incorporating informative samples [25]. In addition, we believe that SSL can also improve morphological explainability by offering more adaptive class definitions. Compared with semi-supervised learning [26], the label-free pretraining stage of SSL models lays a versatile foundation for task-specific fine-tuning that extends beyond cell classification. SSL shows high flexibility by supporting various feature extraction backbones [27]. Among them are lightweight models such as EfficientNet, which have been applied to supervised leukocyte classification [28,29]. Despite advancements like the self-supervised hematology classification model DinoBloom [30], models that integrate morphological insights to clinical workflows through label-efficient leukocyte classification remain largely unexplored. We hypothesize that label-efficient SSL can be extended to support morphological profiling of blood cells, enabling fine-grained differentiation of leukocyte subtypes, particularly blasts [31].

To address the limitations of current automated analyzers and harness SSL, we present a label-efficient approach for leukocyte classification and morphological profiling. Our models were trained and evaluated on over 23,000 single-cell image crops from public and in-house PBS datasets, of which 5,952 in-house images carried hematopathologist-verified cell type labels, including blasts. By applying contrastive learning to unlabeled image triplets, the model generated an embedding space that captured single-cell morphological variations. This morphological landscape facilitated downstream cell type prediction, maintaining high performance even with limited labeled samples. Active learning further boosted label efficiency, revealing cell-type-specific learning efficiency. Moreover, the model was readily fine-tuned for classifying multiple morphological attributes, enhancing explainability. To promote clinical integration, we developed HemoSight, an open-source, containerized web application for interactive PBS review with real-time model inference and interactive morphological landscape visualization.

## Methods

### Data and annotation collection

We obtained blood samples from 43 patients, randomly selected from those admitted to the University of Texas MD Anderson Cancer Center in 2023. Twenty-three patients contributed to more than one sample. Peripheral blood smear (PBS) slides were prepared using Wright’s staining method and scanned using the Sysmex DI-60 automated digital cell morphology system. Approximately 115 white blood cells per slide were sampled using the CellaVision software to generate image crops measuring around 360 × 360 pixels at 1,000× magnification, with each crop centered on a leukocyte. The labels of the image crops were iteratively reviewed by three board-certified hematopathologists (X. H., X. S., Z. H.), using CellaVision classifications as the initial reference, and any disagreements were resolved by consensus.

We obtained additional PBS images from a public dataset [14,32], which consisted of 17,902 images across eight cell type classes: neutrophils, eosinophils, basophils, lymphocytes, monocytes, immature granulocytes (“ig”), erythroblasts, and platelets. The “ig” class includes promyelocytes, myelocytes, and metamyelocytes. Because only healthy individuals were included in the public dataset, we supplemented 5,952 images from our collected samples, resulting in a combined dataset totaling 23,044 images across nine cell type classes. The missing class “blast” from the public dataset was defined to include both blast cells (e.g., myeloblasts, lymphoblasts, and monoblasts) and blast-equivalent cells (e.g., promonocytes). The distribution of classes and cohort demographic statistics are detailed in **Supplementary Tables 1, 2 and 3**.

From the complete dataset, 10% of the images (2,305 crops) were randomly selected using stratified sampling across cell types to form a hold-out test set. The remaining data was split into the training set (∼16,591 images) and the validation set (∼4,148 images) through 5-fold stratified sampling. Grouped splitting by patient or sample was not feasible, as patient-level metadata is unavailable in the public dataset, and many in-house samples contained only a subset of cell types. To further assess patient-level generalization, we collected a new-patient test set comprising 1,109 images from newly acquired slides of 10 distinct patients at the same institution, following identical preparation, staining, scanning, and expert label review procedures (**Supplementary Tables 4 and 5**). This study was approved by the institutional review board at the University of Texas MD Anderson Cancer Center (protocol number 2023-0479-MDACC).

### Label-efficient self-supervised cell typing of leukocytes

We developed a label-efficient framework for leukocyte classification using self-supervised representation learning followed by lightweight supervised fine-tuning. A triplet-based contrastive objective was used for pretext training with unlabeled images, with EfficientNetV2-B0 as the shared encoder backbone initialized from ImageNet weights [33,34]. Input images were preprocessed with color normalization and augmentation before being center-cropped. Anchor-positive-negative triplets of images were derived from augmented views, and training employed semi-hard triplet loss using L2 distance [35]. After pretext training, the encoder weights were frozen, and a linear support vector machine (SVM) was trained on image embeddings to assign cell type labels[36]. For benchmarking, supervised models using the same encoder were trained using labeled images with transfer learning and cross-entropy loss at various labeled data proportions. Model explainability was assessed using Randomized Input Sampling for Explanation (RISE[37]) to generate saliency maps based on randomized masked inputs. All experiments were run in a containerized environment on a high-performance computing cluster equipped with graphics processing units (GPU). Full training, implementation, and deployment details are provided in **Supplementary Methods**.

### Boosting label-efficient learning through active learning

To further enhance label efficiency during the learning process, we applied simulated active learning strategies using the modAL package (version 0.4.2.1) [38] on our dataset with cell-type labels. The process began by initializing the model with an initial batch of *N*_*i*_= 90 labeled samples, randomly selected and stratified across cell types. Subsequent query batches of *N*_*q*_ = 90 samples were drawn from a fixed unlabeled pool. True labels were revealed only for queried samples at each iteration. Sampling was guided by inferred labels and probabilities from the SVM, using two strategies: (a) uncertainty sampling, which prioritized samples with the highest uncertainty (i.e., the lowest maximum probability across all classes), and (b) random sampling, which served as the baseline strategy. It is important to note that this cumulative random sampling approach differs from the random resampling of the training set with increasing sample sizes described in the previous section.

For each iteration, per-class *F*_1_ scores were computed on the validation fold, and the process continued until all training samples had been utilized. To compare the effectiveness of the sampling strategies, we plotted learning curves for each class, representing performance (*P*) as a function of sample size (*n*). These curves were then fitted to an inverse power-law equation (1), where *P*_max_ represents the asymptotic performance, *r* is the learning efficiency, *k* is the scaling factor, and *n*_0_ is the offset.

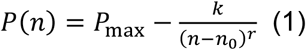

### Mapping the morphological landscape using adaptive fine-tuning on attribute sets

To evaluate the model’s generalization capabilities, we applied it to a subset of the data annotated with 11 morphological attributes sourced from the literature [39]. Since not all images included morphological annotations, we limited the analysis to the intersection of available data points while preserving the original data split. This resulted in approximately 7,379 training images and 1,826 validation images. The classification for each morphological attribute was performed using the same SVM settings.

Uniform Manifold Approximation and Projection (UMAP) [40] was used to visualize the learned morphology landscape by reducing the dimension of embedding vectors from 1280 to 2. The umap-learn package [40] was used with the following hyperparameters configured: number of neighbors set to 15, minimal distance set to 0.1, and Euclidean distance.

To obtain morphological gradients, we performed Linear Discriminant Analysis (LDA) using the sklearn package. Using combined training and validation data, separate one-component LDAs were fitted for each morphological attribute on annotated data and applied to all available data.

## Results

### Label-efficient self-supervised cell typing of leukocytes

To address the need for more efficient workflows compared to the standard analyzers, often inflexible, sensitive, and difficult to interpret, we developed self-supervised models optimized for high label efficiency with minimal performance trade-offs (**Fig. 1**). Our two-stage model begins with a contrastive label-free learning phase followed by fine-tuning using labeled data from a support set (**Fig. S1**). During the contrastive similarity learning phase, image triplets of single-cell crops were generated, consisting of an anchor, a positive, and a negative image. The anchor and positive images were augmentations of the same cell image, while the negative image was drawn from a different cell image. By adjusting the size of the support set, thus the proportion of labeled training data, we effectively lowered the label dependency needed to initialize the training, while enabling further improvement of the model using additional labeled data (**Fig. S1**).

**Fig. 1.**
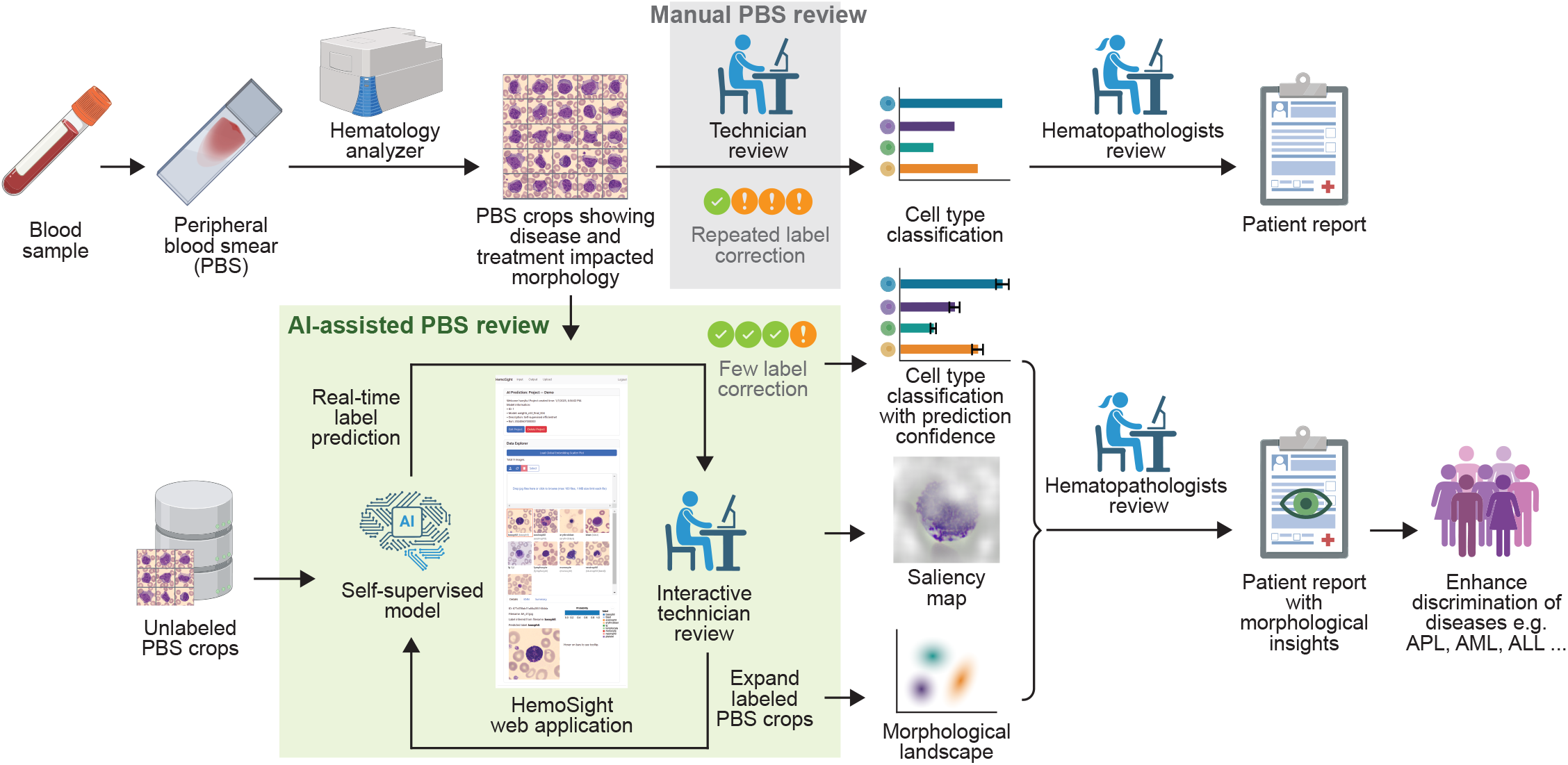
AI-assisted peripheral blood smear review workflow. By leveraging unlabeled images for label efficiency and continuously improving with additional labeled data, the self-supervised model enhances hematopathology workflows: providing prediction confidence, saliency maps, morphological landscape through the interactive web application, HemoSight. These outputs empower hematopathologists with enriched visual insights, supporting more informed and consistent patient reporting, potentially leading to better discrimination of diseases such as acute promyelocytic leukemia (APL), acute myeloid leukemia (AML), and acute lymphoblastic leukemia (ALL)[42].

To evaluate the impact of support set size, we benchmarked our self-supervised models’ performance against supervised models of similar architecture on 9-way cell type classification tasks. The self-supervised models tolerated low proportions of labeled data, with the performance surpassing their supervised counterparts below the inflection point of about 20% of label data used for training (**Fig. 2a**). Note that the decline in performance towards lower label counts, in general, could well be justified by the disproportional amount of labeled data saved. For instance, the *F*_1_ score only dropped by 3.86% with 5.4% of labeled data, from the maximum *F*_1_ score achieved of 96.32%, which required 100% of data to be labeled (**Fig. 2a**). In addition, shifting from a supervised to a self-supervised strategy had minimal impact on the performance ceiling when all labels were provided. Across 5 cross-validation folds, the self-supervised models achieved a maximum *F*_1_ score of 96.32±0.25%, close to 97.47±0.22% of the supervised models (**Fig. 2a**). The performance remained consistent on the hold-out set across 5 repeats (*F*_1_ = 96.20 ± 0.09%) and generalized well to the new-patient test set (*F*_1_ = 83.03 ± 0.40%, **Supplementary Table 4**). These results indicate that self-supervised models offer competitive performance relative to supervised models, especially in label-scarce scenarios.

**Fig. 2.**
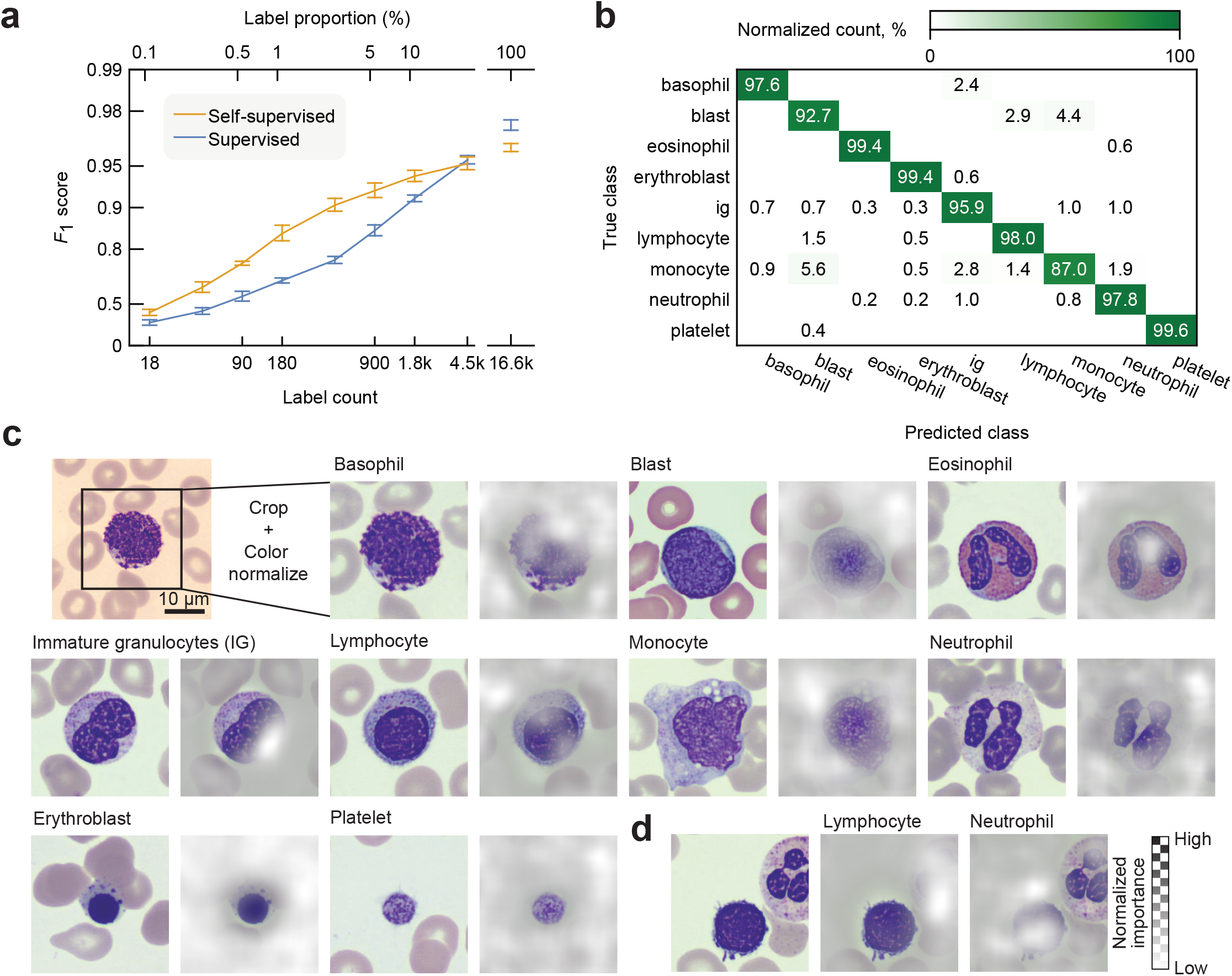
Efficient self-supervised leukocyte typing of peripheral blood smear. (**a**) Cell type prediction performance vs. proportion of labeled data. Values are mean (±SD) across 5-fold cross-validation. The vertical axis is scaled by − log(1 − *x*). (**b**) Confusion matrix shows the performance (one repeat) of the self-supervised model on the hold-out set. Values normalized by count of true class in each class. (**c**) Examples of nine blood cell types relevant to this study next to their respective RISE saliency maps. Top left: inputs to the model are preprocessed on the analyzers’ image crop. (**d**) A representative image crop containing two leukocytes with saliency maps corresponding to two cell type labels. (**c, d**) Saliency maps are range-normalized for each image and to the right of the original image.

We observed that the model performance varies by cell type, attributed to the intrinsic heterogeneity of blood cell morphology. Platelets achieved a near-perfect recall of 99.6%, while monocytes had the lowest recall of 87.0% (**Fig. 2b**). The dominant misclassification occurred between monocytes and blasts, with the highest pairwise error rate of 4.91%. To understand the model decisions across cell types, we employed RISE (Randomized Input Sampling for Explanation [37]) to generate heatmaps highlighting image-based features that were important in the classification task (**Fig. 2c**). High-importance regions can be seen concentrated on the white blood cell rather than the background or surrounding red blood cells, which indicates that the model’s attention falls on center leukocytes and their subcellular features to differentiate cell types, without relying on explicit cell segmentation as expected. Furthermore, in rare cases where images contained more than one leukocyte, the model displayed correct attention on different cells with respect to different labels, demonstrating minimal bias to the location of the cell within the image (**Fig. 2d**).

### Boosting label-efficient learning through active learning

In current clinical settings, label corrections made by hematopathologists are not fed back to the analyzers, leading to repeated classification mistakes. To circumvent this drawback, we hypothesized that active learning sampling strategies could guide incremental expansion of the support set by focusing on informative image-label pairs, thereby maximizing performance gains given the same quantity of new data. The rate of performance improvement with respect to the incremental support set size is hereby referred to as learning efficiency. We performed simulated active learning against ground truth label oracles. We found that the uncertainty sampling strategy, which prioritized inclusion of new samples with the highest uncertainty, drastically reduced the required number of labels. Specifically, only 2,250 labels (13.3% of the total, after 25 iterations) were needed to reach the same performance ceiling as using all labels (**Fig. 3a**). In contrast, random sampling required over 50 batches of images without reaching a clear performance plateau (**Fig. 3a**).

**Fig. 3.**
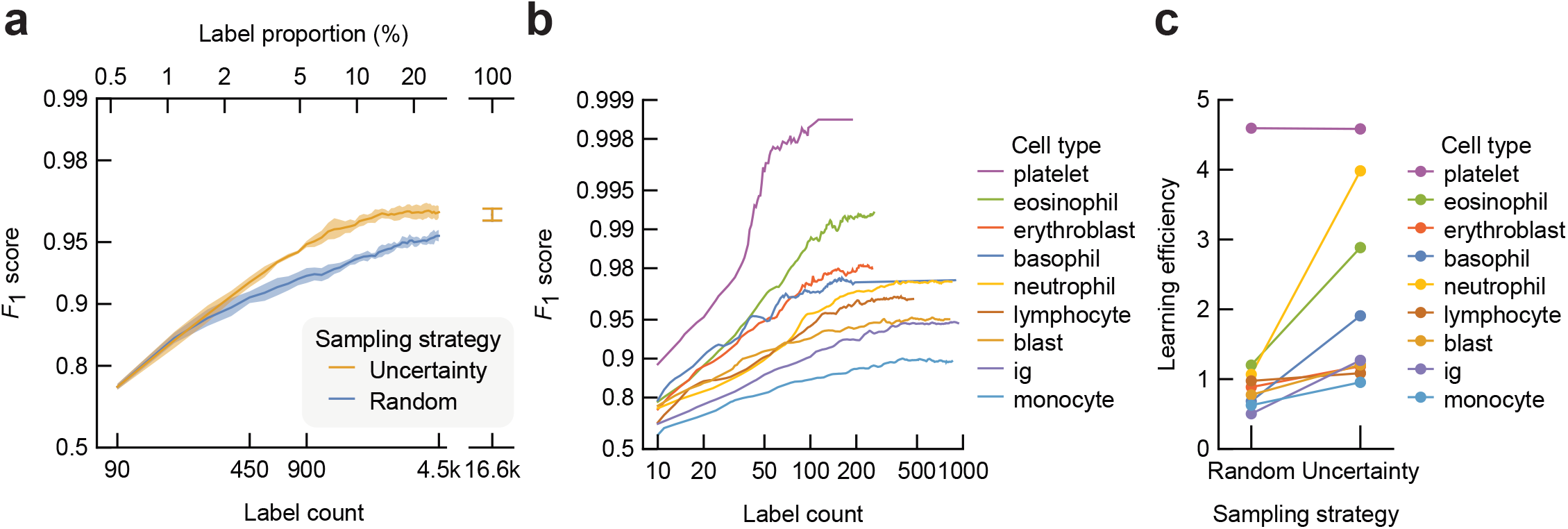
Learning curves vs. active learning sampling strategies. (**a**) Mean macro-averaged *F*_1_ score vs. increasing support set across 5-fold cross-validation. Shaded areas denote SD. (**b**) Learning curves of individual cell types during active learning using uncertainty sampling strategy. Mean values across 5-fold cross-validation are shown. (**c**) Learning efficiency values obtained from mean learning curves of individual cell types from different active learning sampling strategies. (a, b) The vertical axes are scaled by − log(1 − *x*).

In addition, we observed cell-type specific learning efficiency values under the same sampling strategy. Under the uncertainty sampling strategy, among all cell types, the platelet class has the highest learning efficiency, and the monocyte class has the lowest (**Fig. 3b**). We quantified the learning efficiency and the asymptote performance by fitting the learning curve to an inverse power-law function. Per-class learning efficiency values are found to be only weakly correlated to asymptote performance (*ρ* = 0.5, p = 0.170, Spearman’s rank correlation), suggesting the sampling effectiveness and that data quantity alone does not solely determine final performance. Moreover, switching from random to uncertain sampling led to the learning efficiency improvement for 8 of 9 cell types, and the impact of active learning sampling strategies was found to be statistically significant (p = 0.006 one-sided Wilcoxon signed-rank test) (**Fig. 3c**). These results demonstrate that active learning, particularly uncertainty sampling strategy, enhances label efficiency, and that annotation collection should account for cell-type specificity.

### Mapping the morphological landscape using adaptive fine-tuning on attribute sets

We then sought to explore the adaptability of our model by extending its prediction from cell-type labels to morphological features of blood cells. Morphological features offer finer granularity than cell-type annotations and are clinically relevant in PBS review, especially for identifying diseased cell types and detecting rare cells with atypical morphology. Because our pretext training is label-free, we can independently switch label sets and fine-tune the second-stage classifier without reconstructing the embedding space. We obtained 11 distinct morphological attributes of leukocytes from literature [39], encompassing categorical descriptors of shape, color, and texture. The number of categories for each attribute ranges from 2 to 6. Our model demonstrated robust performance, achieving a mean *F*_1_ of 85.78±0.34% across all morphological attributes (**Fig. 4a**). This performance is on par with supervised models (91.20±0.06%) reported in the literature [39]. Notably, the worst-performing attribute was the nucleus shape, which is likely due to its complexity with six categories. Further, we used RISE maps to explain the performance discrepancies (**Fig. 4b**). For the nuclear shape, the model’s attention was scattered rather than focused on the nucleus, whereas it successfully locates regions containing cytoplasm vacuoles. This suggests that it is possible but not guaranteed that the model can identify some subcellular morphological features with only global annotations.

**Fig. 4.**
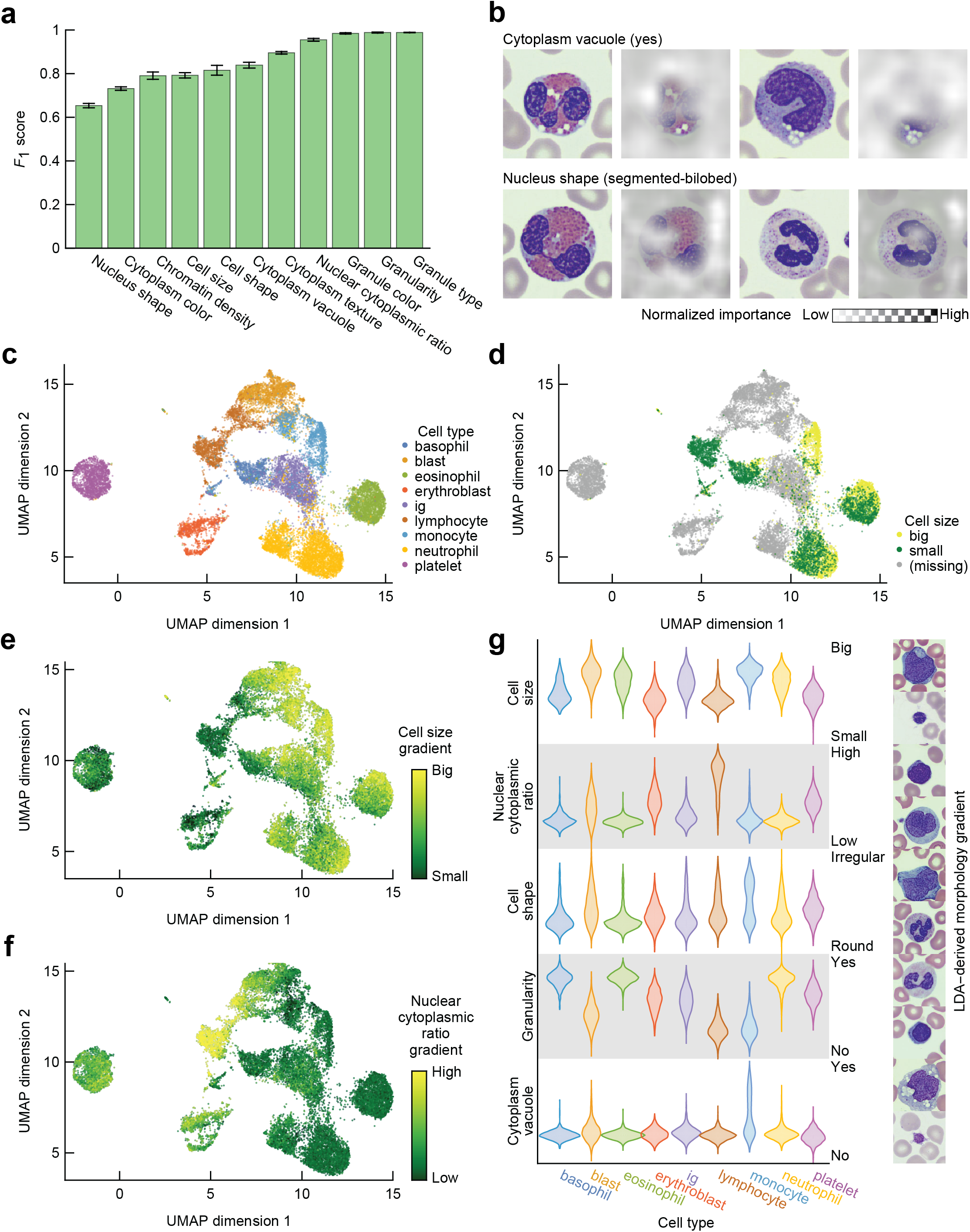
Leukocyte morphological attribute prediction and landscape visualized by UMAP. (**a**) Performance of fine-tuned models across 11 morphological attributes based on learned embedding. Values are mean across 5 folds of cross-validation, and error bars are std. (**b**) Two representative input images and resulting RISE saliency maps from each of two morphological classes. Masked pixels indicate low importance. Saliency maps are range-normalized for each image and to the right of the original image. (**c, d**) Scatter plots showing embedding space dimension reduced by UMAP of a representative model training on the combined training and validation data. Data points are colored by (c) cell type and (d) cell size morphological attributes. (**e, f**) UMAP scatter plots colored by interpolated (e) cell size and (f) nuclear-cytoplasmic ratio gradients obtained from LDA. Colormaps are normalized between 5% to 95% quantile. (**g**) Violin plots showing LDA gradients distributed along various morphological attribute axes for each cell type. Data are range-normalized for each attribute. Sample images correspond to representative cells at the minimum and maximum LDA gradient values for each morphological attribute, selected from non-annotated images.

To visualize model decision boundaries, we reduced the embedding dimensions from 1280 to 2 using Uniform Manifold Approximation and Projection (UMAP). The resulting UMAP embedding (**Fig. 4c**) reveals distinct clusters segregated by cell type. Qualitatively, we note that the large interclass separation may help explain high learning efficiency in some cell types such as platelet and eosinophil (**Fig. 3b**). In contrast, overlapping regions between some cell type clusters are consistent with the error cases shown in the confusion matrix (**Fig. 2b**). When projecting morphological features, such as cell size, onto the UMAP embedding, we observed polarized distributions within certain cell types, including eosinophils and neutrophils (**Fig. 4d**). This suggests that the learned morphological landscape captures similarities driven by both cell type distinctions and underlying morphological characteristics.

Finally, we performed linear discriminant analysis (LDA) on the original feature vectors to obtain the best separation of categories of each morphological attribute. The solutions of LDA enable disentanglement of the morphological landscape along individual attributes and interpolation of morphological gradients from discretized categories to continuous variables, such as cell size (**Fig. 4e**) and nucleus-to-cytoplasm (N/C) ratio (**Fig. 4f**). Since not all images had morphological annotations, and some morphological attributes were not applicable to certain cell types (e.g. anucleate platelet), LDA enables us to orientate within the morphological landscape and probe model’s learning of relative cytomorphology similarities. Compared to **Fig. 4d, Fig. 4e** gives a fuller picture of cell size distributions, which correctly position platelets and erythroblasts towards relatively small cell sizes. From **Fig. 4f**, we can see that blasts similar to lymphocytes have a high N/C ratio, while those similar to monocytes have a low N/C ratio, indicating a sub-cluster distribution that aligns with possible blast subtypes. By examining LDA-derived morphological gradients (**Fig. 4g**), we may have more qualitative observations reflecting cell-type-specific morphological features. For example, cytoplasm vacuoles occur more often in monocytes; cell shape irregularity is more prominent in monocytes and blasts. Overall, the learned embeddings demonstrate strong predictive power, effectively generalizing to categorize morphological attributes in unseen cases with limited annotations.

### Real-time model inferencing through HemoSight web application

Finally, inspired by the challenge of creating an open platform for seamless cell classification to support medical image analysis [41], we developed HemoSight, a browser-accessible implementation of our self-supervised leukocyte classification model. HemoSight allows end users to batch-upload images and obtain cell-type predictions in real time (**Fig. 1, Fig. S2a**), eliminating the need for local computational resources capable of running deep-learning models. HemoSight also includes the confidence of classification, which could help to make informed decisions based on the system’s output uncertainty (**Fig. S2a**). Feedback from the three board-certified hematopathologists involved in this study was incorporated to design a user interface that reflects the workflows of commercial software and to include classification confidence scores to facilitate manual label correction.

HemoSight is a full-stack application that is containerized and deployed on a server (**Fig. S2b**). By offloading computations to the server and leveraging a lightweight feature encoder, the application achieves near-instantaneous processing, delivering tens of predictions per second without parallel processing optimizations. Beyond classification, HemoSight provides informative visualizations not available in conventional analyzers, such as prediction confidence levels and similarity-based image retrieval (**Fig. S2c**). This web application lays the foundation for AI-assisted hematopathology workflows, enhancing PBS review with integrated image analysis, visual interpretation, and quantitative morphological measures in patient reports.

## Discussion

In this study, we presented an annotation-efficient and explainable leukocyte classification framework designed to enhance peripheral blood smear (PBS) review in clinical laboratories. We curated a dataset that includes cell types such as blasts, which are crucial for diagnostic decisions in hematological malignancies. Central to this framework is a self-supervised deep learning model that accommodates flexible label sets, such as cell types and morphological features. Our self-supervised model demonstrated strong resilience to label scarcity benchmarked against supervised approaches, and the label efficiency was further enhanced through active learning. Our framework provided critical explainability and confidence for model decisions, which prompted us to integrate the model into HemoSight, an interactive web application.

Our study has the potential to generate transformative clinical utility for AI-assisted hematopathology workflows. We envision that next-generation patient reports will integrate AI-driven cytomorphology analysis to bolster consistency and efficiency in blood disorder assessment (**Fig. 1**). By providing standardized and reproducible morphological insights, AI can assist hematopathologists in differentiating overlapping hematologic conditions, minimizing diagnostic variability, and improving early detection of aggressive diseases [42]. For example, this capability has immediate applications in refining the classification of abnormal lymphocytes in leukemia and lymphoma subtypes [43], differentiating non-Hodgkin lymphoma from acute leukemia [44], and improving recognition of rare leukemia subtypes such as plasma cell leukemia [45] and acute erythroid leukemia [46]. In addition, longitudinal studies across patient cohorts could uncover subtle, disease-driven morphological shifts, deepening our understanding of hematologic progression and treatment response [47]. As full-field PBS scanners become more widespread [48], our approach is well-positioned to drive the development of AI-assisted grading of whole slide images, enabling more scalable and automated workflows. Beyond PBS, our framework could be adapted for bone marrow biopsy interpretation [49,50] or imaging flow cytometry [51], with tailored roadmaps for model development and clinical integration, as demonstrated by recent AI-assisted digital microscopy systems [52]. These would involve standardizing sample preparation and digitization, implementing additional preprocessing such as cell detection and segmentation, establishing cohorts with well-represented cases and annotated subsets, exploring clinically compatible reporting formats and interpretability feedback, conducting prospective evaluation, and addressing regulatory and infrastructure requirements for deployment. To sum up, after extensive self-supervised training on label-free datasets across one or more domains, our workflow builds a strong foundation for label-efficient fine-tuning, readily extending to various domain-specific classification tasks.

Despite promising results, our approach presents several limitations and opportunities for future work. Firstly, we did not assess the model’s ability to generalize over local features. The feature embedding is derived from the entire image crops centered around the leukocyte, and the morphological attributes are mainly defined by global features, except for the cytoplasm vacuole. Subcellular morphologies, e.g. Auer rods, Döhle bodies, hairy-like cytoplasmic projections [10], are crucial for diagnosing certain hematological disorders. To prevent local features from being overwhelmed by global ones in the learned feature representation, future work could explore alternative image augmentation strategies or model architectures designed to better capture local features, such as multi-scale convolutional neural networks (CNN) [53] or transformer-based models with local-global attention mechanisms [54]. Alternatively, large-scale foundation models such as DinoBloom [30] may serve as plug-in encoders for more robust morphological features, but their computational cost would need to be carefully monitored in latency-sensitive, real-time applications. Secondly, additional datasets are necessary to evaluate batch differences and to further validate the model. While color normalization was implemented to account for systemic variations, extending the dataset to include a broader patient population and multiple institutions, alongside patient-level splitting, could enhance the robustness of the learned morphological landscape and improve prediction accuracy. Thirdly, exploring alternative fine-tuning classifiers and active learning strategies could further improve model performance. We opted for a support vector machine (SVM) to balance simplicity and performance, but alternative classifiers, such as tree-based ensemble methods, could be tested, with careful monitoring of overfitting and computational overhead. Likewise, we could explore additional active learning sampling strategies, such as density-based, representativeness-based or hybrid sampling [55], and evaluate the impact of initialization techniques [56] or query batch size on performance. Finally, although the HemoSight web application demonstrated real-time model inference, it remains a work in progress. While supported by our self-supervised learning framework, the application does not yet incorporate label correction feedback or dynamic class redefinition—key features for iterative refinement of predicted leukocyte labels, which are essential to implement active learning beyond oracle-based labeling.

In conclusion, our work represents a pioneering effort in advancing label-efficient, morphology-aware AI models for hematopathology. By fostering a synergistic relationship between AI and hematopathologists, we aim to accelerate the seamless integration of adaptive AI systems into clinical practice, ultimately enhancing the accuracy, efficiency, and reproducibility of PBS review in patient care.

## Contributors

Z.L. developed methods, performed model comparisons, and generated and visualized results. Z.L. developed and deployed the web application. Y.Y. and S.P.C. supervised and provided guidance on method development, comparison, and analysis. Z.H. collected and annotated in-house PBS crops, which were reviewed by X.S. and X.H. Z.H., X.S., and X.H. further provided clinical insight. Z.H. prepared and submitted the IRB review for the study. Z.L., S.P.C. and Y.Y. wrote the manuscript with input from Z.H., X.S.. Z.H. and Y.Y. conceived the research.

## Declaration of Interests

The authors declare no potential conflicts of interest.

## Acknowledgment

We thank Xiaoxi Pan for her expertise and discussion in model training. This work is supported by seed funding from the University of Texas MD Anderson Cancer Center to Y.Y. Generative AI was used to refine the text in the manuscript with all edits manually reviewed for clarity and accuracy.

## Data Sharing Statements

Data available upon request to Dr. Zhihong Hu.

## Notes

### Competing Interest Statement

The authors have declared no competing interest.

## References

[1] Gulati G, Song J, Florea AD, Gong J. Purpose and criteria for blood smear scan, blood smear examination, and blood smear review. Ann Lab Med 2013;33:1–7. 10.3343/ALM.2013.33.1.1.

[2] Zini G. Abnormalities in leukocyte morphology and number. Blood and Bone Marrow Pathology, Elsevier; 2011, p. 247–61. 10.1016/B978-0-7020-3147-2.00016-X.

[3] Bates I, Bain BJ. Approach to the diagnosis and classification of blood diseases. Dacie and Lewis Practical Haematology 2012:549. 10.1016/B978-0-7020-3408-4.00023-0.

[4] Matutes E, Polliack A. Morphological and Immunophenotypic Features of Chronic Lymphocytic Leukemia. Rev Clin Exp Hematol 2000;4:22–47. 10.1046/J.1468-0734.2000.00002.X.

[5] Pratumvinit B, Wongkrajang P, Reesukumal K, Klinbua C, Niamjoy P. Validation and Optimization of Criteria for Manual Smear Review Following Automated Blood Cell Analysis in a Large University Hospital. Arch Pathol Lab Med 2013;137:408–14. 10.5858/ARPA.2011-0535-OA.

[6] Coombs CC, Tavakkoli M, Tallman MS. Acute promyelocytic leukemia: where did we start, where are we now, and the future. Blood Cancer Journal 2015 5:4 2015;5:e304– e304. 10.1038/bcj.2015.25.

[7] Kratz A, Bengtsson HI, Casey JE, Keefe JM, Beatrice GH, Grzybek DY, et al. Performance Evaluation of the CellaVision DM96 System: WBC Differentials by Automated Digital Image Analysis Supported by an Artificial Neural Network. Am J Clin Pathol 2005;124:770–81. 10.1309/XMB9K0J41LHLATAY.

[8] Gulati G, Uppal G, Gong J. Unreliable Automated Complete Blood Count Results: Causes, Recognition, and Resolution. Ann Lab Med 2022;42:515–30. 10.3343/ALM.2022.42.5.515.

[9] Wang X, Wang X, Ge P, Zhao X, Chen C, Hu S, et al. Establishment of improved review criteria for hematology analyzers in cancer hospitals. J Clin Lab Anal 2021;35. 10.1002/JCLA.23638.

[10] Hegde RB, Prasad K, Hebbar H, Sandhya I. Peripheral blood smear analysis using image processing approach for diagnostic purposes: A review. Biocybern Biomed Eng 2018;38:467–80. 10.1016/J.BBE.2018.03.002.

[11] Kratz A, Lee S hee, Zini G, Riedl JA, Hur M, Machin S. Digital morphology analyzers in hematology: ICSH review and recommendations. Int J Lab Hematol 2019;41:437–47. 10.1111/ijlh.13042.

[12] Suganya Devi K, Arutperumjothi G, Srinivasan P. Diagnosis Evaluation and Interpretation of Qualitative Abnormalities in Peripheral Blood Smear Images—A Review. Studies in Computational Intelligence 2021;932:341–65. 10.1007/978-981-15-9735-0_17.

[13] Gedefaw L, Liu CF, Ip RKL, Tse HF, Yeung MHY, Yip SP, et al. Artificial Intelligence-Assisted Diagnostic Cytology and Genomic Testing for Hematologic Disorders. Cells 2023, Vol 12, Page 1755 2023;12:1755. 10.3390/CELLS12131755.

[14] Acevedo A, Alférez S, Merino A, Puigví L, Rodellar J. Recognition of peripheral blood cell images using convolutional neural networks. Comput Methods Programs Biomed 2019;180:105020. 10.1016/J.CMPB.2019.105020.

[15] Chola C, Muaad AY, Bin Heyat MB, Benifa JVB, Naji WR, Hemachandran K, et al. BCNet: A Deep Learning Computer-Aided Diagnosis Framework for Human Peripheral Blood Cell Identification. Diagnostics 2022, Vol 12, Page 2815 2022;12:2815. 10.3390/DIAGNOSTICS12112815.

[16] Anand V, Gupta S, Koundal D, Alghamdi WY, Alsharbi BM. Deep learning-based image annotation for leukocyte segmentation and classification of blood cell morphology. BMC Med Imaging 2024;24:1–15. 10.1186/S12880-024-01254-Z/TABLES/3.

[17] Lynch DT, Hall J, Foucar K. How I investigate monocytosis. Int J Lab Hematol 2018;40:107–14. 10.1111/IJLH.12776.

[18] Wang Q, Bi S, Sun M, Wang Y, Wang D, Yang S. Deep learning approach to peripheral leukocyte recognition. PLoS One 2019;14:e0218808. 10.1371/JOURNAL.PONE.0218808.

[19] Lee LH, Mansoor A, Wood B, Nelson H, Higa D, Naugler C. Performance of CellaVision DM96 in leukocyte classification. J Pathol Inform 2013;4:14. 10.4103/2153-3539.114205.

[20] Kim HN, Hur M, Kim H, Kim SW, Moon HW, Yun YM. Performance of automated digital cell imaging analyzer Sysmex DI-60. Clin Chem Lab Med 2017;56:94–102. 10.1515/CCLM-2017-0132/MACHINEREADABLECITATION/RIS.

[21] De Iuliis V, Chiatamone Ranieri S. Performance Evaluation of the Scopio Labs X100HT Digital Morphology Analyzer and Abnormal Cell Detection in Peripheral Blood Smears. Int J Lab Hematol 2025;0:1–9. 10.1111/IJLH.70005.

[22] Huang SC, Pareek A, Jensen M, Lungren MP, Yeung S, Chaudhari AS. Self-supervised learning for medical image classification: a systematic review and implementation guidelines. Npj Digital Medicine 2023 6:1 2023;6:1–16. 10.1038/s41746-023-00811-0.

[23] Nielsen M, Wenderoth L, Sentker T, Werner R. Self-Supervision for Medical Image Classification: State-of-the-Art Performance with ∼100 Labeled Training Samples per Class. Bioengineering 2023;10. 10.3390/BIOENGINEERING10080895.

[24] Khan S, Sajjad M, Hussain T, Ullah A, Imran AS. A review on traditional machine learning and deep learning models for WBCs classification in blood smear images. IEEE Access 2021;9:10657–73. 10.1109/ACCESS.2020.3048172.

[25] Bengar JZ, van de Weijer J, Twardowski B, Raducanu B. Reducing Label Effort: Self-Supervised Meets Active Learning 2021:1631–9.

[26] Song H, Wang Z. Automatic Classification of White Blood Cells Using a Semi-Supervised Convolutional Neural Network. IEEE Access 2024;12:44972–83. 10.1109/ACCESS.2024.3380896.

[27] Ornob TR, Roy G, Hassan E. CovidExpert: A Triplet Siamese Neural Network framework for the detection of COVID-19. Inform Med Unlocked 2023;37:101156. 10.1016/J.IMU.2022.101156.

[28] Batool A, Byun YC. Lightweight EfficientNetB3 Model Based on Depthwise Separable Convolutions for Enhancing Classification of Leukemia White Blood Cell Images. IEEE Access 2023;11:37203–15. 10.1109/ACCESS.2023.3266511.

[29] Teoh TT. CNN for White Blood Cell Classification. SpringerBriefs in Computer Science 2023:53–68. 10.1007/978-981-19-8814-1_4/FIGURES/9.

[30] Koch V, Wagner SJ, Kazeminia S, Sancar E, Hehr M, Schnabel JA, et al. DinoBloom: A Foundation Model for Generalizable Cell Embeddings in Hematology 2024:520–30. 10.1007/978-3-031-72390-2_49.

[31] Dinčić M, Popović TB, Kojadinović M, Trbovich AM, Ilić A. Morphological, fractal, and textural features for the blood cell classification: the case of acute myeloid leukemia. European Biophysics Journal 2021;50:1111–27. 10.1007/S00249-021-01574-W/FIGURES/10.

[32] Acevedo A, Merino A, Alférez S, Molina Á, Boldú L, Rodellar J. A dataset of microscopic peripheral blood cell images for development of automatic recognition systems. Data Brief 2020;30:105474. 10.1016/J.DIB.2020.105474.

[33] Tan M, Le Q V. EfficientNetV2: Smaller Models and Faster Training. Proc Mach Learn Res 2021;139:10096–106.

[34] Deng J, Dong W, Socher R, Li L-J, Kai Li, Li Fei-Fei. ImageNet: A large-scale hierarchical image database 2010:248–55. 10.1109/CVPR.2009.5206848.

[35] Schroff F, Kalenichenko D, Philbin J. FaceNet: A Unified Embedding for Face Recognition and Clustering. Proceedings of the IEEE Computer Society Conference on Computer Vision and Pattern Recognition 2015;07-12-June-2015:815–23. 10.1109/cvpr.2015.7298682.

[36] Cortes C, Vapnik V, Saitta L. Support-vector networks. Machine Learning 1995 20:3 1995;20:273–97. 10.1007/BF00994018.

[37] Petsiuk V, Das A, Saenko K. RISE: Randomized Input Sampling for Explanation of Black-box Models. British Machine Vision Conference 2018, BMVC 2018 2018.

[38] Danka T, Horvath P. modAL: A modular active learning framework for Python 2018.

[39] Tsutsui S, Pang W, Wen B. WBCAtt: A White Blood Cell Dataset Annotated with Detailed Morphological Attributes. Adv Neural Inf Process Syst 2023;36:50796–824.

[40] McInnes L, Healy J, Melville J. UMAP: Uniform Manifold Approximation and Projection for Dimension Reduction 2018.

[41] Escobar Díaz Guerrero R, Carvalho L, Bocklitz T, Popp J, Oliveira JL. Software tools and platforms in Digital Pathology: a review for clinicians and computer scientists. J Pathol Inform 2022;13:100103. 10.1016/J.JPI.2022.100103.

[42] Chase ML, Drews R, Zumberg MS, Ellis LR, Reid EG, Gerds AT, et al. Consensus recommendations on peripheral blood smear review: defining curricular standards and fellow competency. Blood Adv 2023;7:3244–52. 10.1182/BLOODADVANCES.2023009843.

[43] Rodellar J, Alférez S, Acevedo A, Molina A, Merino A. Image processing and machine learning in the morphological analysis of blood cells. Int J Lab Hematol 2018;40:46–53. 10.1111/IJLH.12818.

[44] Dorfman DM, Sadigh S. Non-Hodgkin lymphoma mimicking acute leukemia: a report of six cases and review of the literature. J Hematop 2022;15:63. 10.1007/S12308-022-00493-9.

[45] Sekar MD, Raj M, Manivannan P. Role of Morphology in the Diagnosis of Acute Leukemias: Systematic Review. Indian Journal of Medical and Paediatric Oncology 2023;44:464–73. 10.1055/S-0043-1764369/ID/JR222811059-42/BIB.

[46] Reichard KK, Tefferi A, Abdelmagid M, Orazi A, Alexandres C, Haack J, et al. Pure (acute) erythroid leukemia: morphology, immunophenotype, cytogenetics, mutations, treatment details, and survival data among 41 Mayo Clinic cases. Blood Cancer J 2022;12:1–8. 10.1038/S41408-022-00746-X;TECHMETA=45;SUBJMETA=1541,1897,1990,283,631,67,692,699;KWRD=ACUTE+MYELOID+LEUKAEMIA.

[47] Chabot-Richards DS, Foucar K. Does morphology matter in 2017? An approach to morphologic clues in non-neoplastic blood and bone marrow disorders. Int J Lab Hematol 2017;39:23–30. 10.1111/IJLH.12667.

[48] Katz BZ, Feldman MD, Tessema M, Benisty D, Toles GS, Andre A, et al. Evaluation of Scopio Labs X100 Full Field PBS: The first high-resolution full field viewing of peripheral blood specimens combined with artificial intelligence-based morphological analysis. Int J Lab Hematol 2021;43:1408–16. 10.1111/IJLH.13681.

[49] Eckardt JN, Schmittmann T, Riechert S, Kramer M, Sulaiman AS, Sockel K, et al. Deep learning identifies Acute Promyelocytic Leukemia in bone marrow smears. BMC Cancer 2022;22:201. 10.1186/S12885-022-09307-8.

[50] Tripathi S, Augustin AI, Sukumaran R, Dheer S, Kim E. HematoNet: Expert level classification of bone marrow cytology morphology in hematological malignancy with deep learning. Artificial Intelligence in the Life Sciences 2022;2:100043. 10.1016/J.AILSCI.2022.100043.

[51] Lippeveld M, Knill C, Ladlow E, Fuller A, Michaelis LJ, Saeys Y, et al. Classification of Human White Blood Cells Using Machine Learning for Stain-Free Imaging Flow Cytometry. Cytometry Part A 2020;97:308–19. 10.1002/CYTO.A.23920.

[52] Bagg A, Raess PW, Rund D, Bhattacharyya S, Wiszniewska J, Horowitz A, et al. Performance Evaluation of a Novel Artificial Intelligence–Assisted Digital Microscopy System for the Routine Analysis of Bone Marrow Aspirates. Modern Pathology 2024;37:100542. 10.1016/J.MODPAT.2024.100542.

[53] Huo X, Sun G, Tian S, Wang Y, Yu L, Long J, et al. HiFuse: Hierarchical multi-scale feature fusion network for medical image classification. Biomed Signal Process Control 2024;87:105534. 10.1016/J.BSPC.2023.105534.

[54] Oquab M, Darcet T, Moutakanni T, Vo H V., Szafraniec M, Khalidov V, et al. DINOv2: Learning Robust Visual Features without Supervision. Transactions on Machine Learning Research 2023;2024.

[55] Wang H, Jin Q, Li S, Liu S, Wang M, Song Z. A comprehensive survey on deep active learning in medical image analysis. Med Image Anal 2024;95:103201. 10.1016/J.MEDIA.2024.103201.

[56] Yang Y, Loog M. To Actively Initialize Active Learning. Pattern Recognit 2022;131:108836. 10.1016/J.PATCOG.2022.108836.

[57] Ioffe S, Szegedy C. Batch Normalization: Accelerating Deep Network Training by Reducing Internal Covariate Shift. 32nd International Conference on Machine Learning, ICML 2015 2015;1:448–56.

[58] Buchsbaum G. A spatial processor model for object colour perception. J Franklin Inst 1980;310:1–26. 10.1016/0016-0032(80)90058-7.

[59] Satyanarayan A, Moritz D, Wongsuphasawat K, Heer J. Vega-Lite: A Grammar of Interactive Graphics. IEEE Trans Vis Comput Graph 2017;23:341–50. 10.1109/TVCG.2016.2599030.

