## Supplementary Materials for "Interpretable morphology mapping of peripheral blood leukocytes using annotation-efficient artificial intelligence"

### Supplementary Figures

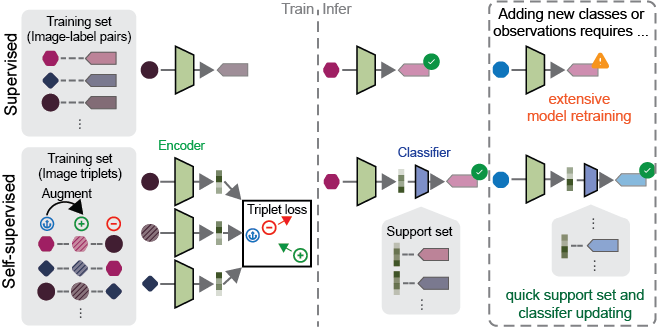

Fig. S1 Adaptability of self-supervised vs. supervised classification models to new classes or observations. By decoupling label information to the support set, the self-supervised approach enables rapid adaptation to newly defined classes or additional samples after training, unlike traditional supervised models, which require lengthy retraining.

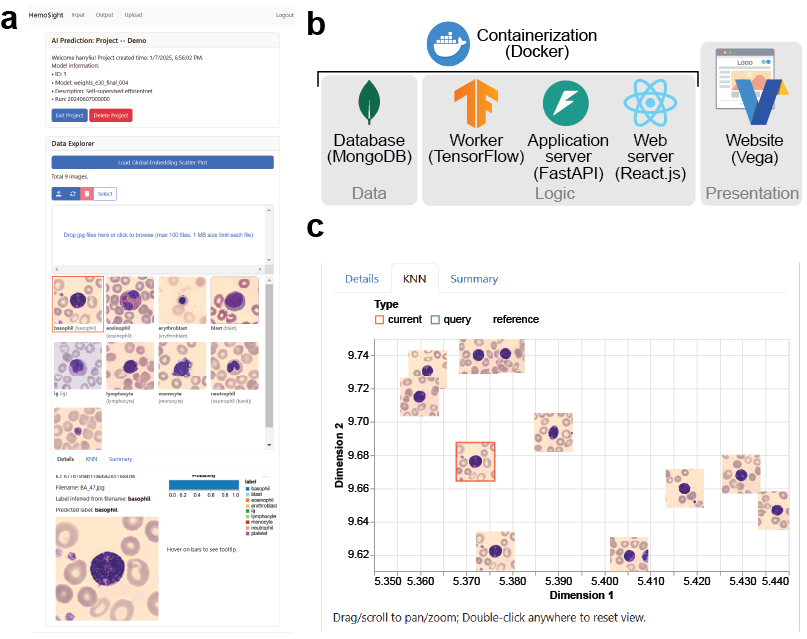

Fig. S2 Construction and features of the HemoSight web application. (**a**) HemoSight enables users to upload query images and perform cell type inference. The prediction uncertainty for each image is shown at the bottom of the page in the currently displayed tab. (**b**) Frameworks and tools that enabled the HemoSight web application. (**c**) Given one query image, its nearest neighbors can be visualized in the embedding space which is dimensionally reduced by UMAP.

### Supplementary Table

| Class | Public dataset image count | In-house dataset image count | Total count |
| --- | --- | --- | --- |
| basophil | 1218 (7.1%) | 14 (0.2%) | 1232 (5.3%) |
| blast | 0 (0.0%) | 2738 (46.0%) | 2738 (11.9%) |
| eosinophil | 3117 (18.2%) | 62 (1.0%) | 3179 (13.8%) |
| erythroblast | 1551 (9.1%) | 0 (0.0%) | 1551 (6.7%) |
| “ig” | 2895 (16.9%) | 18 (0.3%) | 2913 (12.6%) |
| lymphocyte | 1214 (7.1%) | 800 (13.4%) | 2014 (8.7%) |
| monocyte | 1420 (8.3%) | 724 (12.2%) | 2144 (9.3%) |
| neutrophil | 3329 (19.5%) | 1596 (26.8%) | 4925 (21.4%) |
| platelet | 2348 (13.7%) | 0 (0.0%) | 2348 (10.2%) |
| Total | 17092 | 5952 | 23044 |

**Table S1 Distribution of images used in this study by cell type classes**. Column-wise proportion shown as percentages in parentheses. The public dataset is from [1]. “ig”: immature granulocytes, which include promyelocytes, myelocytes, and metamyelocytes.

| *Feature* | *Class* | *Count* | *Percentage* |
| --- | --- | --- | --- |
| *Cell type* | *basophil* | 1218 | 11.8% |
|  | *blast* | 0 | 0.0% |
|  | *eosinophil* | 3117 | 30.3% |
|  | *erythroblast* | 1551 | 15.1% |
|  | *“ig”* | 2895 | 28.1% |
|  | *lymphocyte* | 1214 | 11.8% |
|  | *monocyte* | 1420 | 13.8% |
|  | *neutrophil* | 3329 | 32.3% |
|  | *platelet* | 2348 | 22.8% |
| *Cell size* | *big* | 5561 | 54.0% |
|  | *small* | 4737 | 46.0% |
| *Cell shape* | *irregular* | 2332 | 22.6% |
|  | *round* | 7966 | 77.4% |
| *Nucleus shape* | *irregular* | 870 | 8.4% |
|  | *segmented-bilobed* | 3119 | 30.3% |
|  | *segmented-multilobed* | 1284 | 12.5% |
|  | *unsegmented-band* | 2614 | 25.4% |
|  | *unsegmented-indented* | 1342 | 13.0% |
|  | *unsegmented-round* | 1069 | 10.4% |
| *Nuclear cytoplasmic ratio* | *high* | 1238 | 12.0% |
|  | *low* | 9060 | 88.0% |
| *Chromatin density* | *densely* | 9369 | 91.0% |
|  | *loosely* | 929 | 9.0% |
| *Cytoplasm vacuole* | *no* | 9509 | 92.3% |
|  | *yes* | 789 | 7.7% |
| *Cytoplasm texture* | *clear* | 8254 | 80.2% |
|  | *frosted* | 2044 | 19.8% |
| *Cytoplasm color* | *blue* | 1402 | 13.6% |
|  | *light blue* | 7801 | 75.8% |
|  | *purple blue* | 1095 | 10.6% |
| *Granule type* | *coarse* | 1216 | 11.8% |
|  | *nil* | 2631 | 25.5% |
|  | *round* | 3115 | 30.2% |
|  | *small* | 3336 | 32.4% |
| *Granule color* | *nil* | 2630 | 25.5% |
|  | *pink* | 3245 | 31.5% |
|  | *purple* | 1305 | 12.7% |
|  | *red* | 3118 | 30.3% |
| *Granularity* | *no* | 2629 | 25.5% |
|  | *yes* | 7669 | 74.5% |
| *Total* |  | *10298* | *100.0%* |

**Table S2 Distribution of images used in this study by morphological attributes.** Note that morphological labels were only available for the public dataset [1]. “ig”: immature granulocytes, which include promyelocytes, myelocytes, and metamyelocytes.

| *Characteristics* | *Category* | *Count* | *Percentage* |
| --- | --- | --- | --- |
| Sex | *Female* | 15 | 35% |
|  | *Male* | 28 | 65% |
| *Age* | *0-17* | 1 | 2% |
|  | *18-44* | 12 | 28% |
|  | *45-64* | 6 | 14% |
|  | *65-79* | 18 | 42% |
|  | *80+* | 6 | 14% |
| *Diagnosis* | *Acute myeloid leukemia* | 26 | 60% |
|  | *Acute lymphoblastic leukemia* | 5 | 12% |
|  | *Mixed phenotype acute leukemia* | 1 | 2% |
|  | *Chronic myelomonocytic leukemia* | 3 | 7% |
|  | *Reactive monocytosis following therapy* | 8 | 19% |
| *Total* |  | *43* | *100%* |

**Table S3 Patient cohort demographic statistics.**

| Class | Hold-out test set $F_{1}$ score | Hold-out test set image count (n = 2304) | New-patient test set $F_{1}$ score | New-patient test set image count (n = 1109) |
| --- | --- | --- | --- | --- |
| basophil | 97.24±0.33% | 123 (5.3%) | 91.77±1.18% | 54 (4.9%) |
| blast | 93.38±0.31% | 274 (11.9%) | 83.90±0.36% | 117 (10.6%) |
| eosinophil | 99.15±0.18% | 318 (13.8%) | 94.44±0.98% | 57 (5.1%) |
| erythroblast | 98.15±0.41% | 155 (6.7%) | 96.35±0.79% | 37 (3.3%) |
| “ig” | 95.42±0.33% | 291 (12.6%) | 22.82±2.69% | 23 (2.1%) |
| lymphocyte | 95.88±0.41% | 201 (8.7%) | 93.84±0.41% | 200 (18.0%) |
| monocyte | 88.91±0.33% | 215 (9.3%) | 84.25±0.40% | 126 (11.4%) |
| neutrophil | 97.87±0.13% | 493 (21.4%) | 97.23±0.16% | 467 (42.1%) |
| platelet | 99.79±0.00% | 235 (10.2%) | 82.71±4.52% | 28 (2.5%) |
| Macro-averaged $F_{1}$ | 96.20±0.09% |  | 83.03±0.40% |  |

**Table S4** $\boldsymbol{F}_{\boldsymbol{1}}$ **scores and counts of test sets**. For counts, column-wise proportion shown as percentages in parentheses. For $F_{1}$ scores, mean and standard deviation across 5 repeats are shown. “ig”: immature granulocytes, which include promyelocytes, myelocytes, and metamyelocytes.

| *Characteristics* | *Category* | *Count* | *Percentage* |
| --- | --- | --- | --- |
| *Sex* | *Female* | 4 | 40% |
|  | *Male* | 6 | 60% |
| *Age* | *0-17* | 0 | 0% |
|  | *18-44* | 1 | 10% |
|  | *45-64* | 4 | 40% |
|  | *65-79* | 5 | 50% |
|  | *80+* | 0 | 0% |
| *Diagnosis* | *Acute myeloid leukemia* | 4 | 40% |
|  | *Plasma cell neoplasm* | 2 | 20% |
|  | *Myelodysplastic/myeloproliferative neoplasm* | 2 | 20% |
|  | *Chronic myeloid leukemia* | 1 | 10% |
|  | *Reactive monocytosis following therapy* | 1 | 10% |
| *Total* |  | *10* | *100%* |

**Table S5 Patient cohort demographic statistics of the new-patient test set.**

### Supplementary Methods

#### Self-Supervised Pretext Training and Supervised Fine-Tuning

##### Label-free pretext training

We implemented the model using TensorFlow 2.10.1 and Python 3.8.10. Our self-supervised model features three encoder heads with shared weights, followed by a similarity layer. EfficientNetV2-B0 was selected as the feature encoder for its optimized size, which reduces computational costs and accelerates processing [2]. The model was initialized with weights pre-trained on ImageNet [3], and all layers were trainable, except batch normalization layers were frozen [4].

Input images underwent preprocessing steps, including color normalization using the gray world algorithm [5], followed by image augmentation. Augmentation methods included random horizontal and vertical flipping, rotation (±90°), zoom (±0.1×), horizontal and vertical translation (±20 pixels), and color jitter (brightness, contrast, saturation, and hue with a magnitude of 0.2). Images were then center-cropped to 224 × 224 pixels before being fed into the encoder, with pixel intensity rescaling from 0–255 to 0–1 handled internally by TensorFlow.

The output feature vectors from the encoder’s top activation layer were passed through a global average pooling layer, followed by a dropout layer with a rate of 0.2. These vectors were subsequently $L_{2}$-normalized via a lambda layer before loss computation. The pretext similarity-based training relied on anchor-positive-negative triplets, where the anchor and positive images were derived from augmentations of the same cell image, while negative images were selected from different cell images. Triplet semi-hard loss [6] was used, which was implemented by the TensorFlow Addons library (version 0.20). To enable efficient online triplet mining, the image generator produced two augmented versions of each image in a batch so that no triplets were arranged explicitly. The pretext training utilized semi-hard loss with a margin of 0.5 and the $L_{2}$ distance metric. The model was trained for 30 epochs with a batch size of 100 using the Adam optimizer. The learning rate was set to ${10}^{-5}$ with default optimizer parameters of $\beta_{1}=0.9$, $\beta_{2}=0.999$, and $\epsilon={10}^{-7}$. We manually explored the hyperparameter space, including the choice of optimizer, learning rate, scheduling strategies, and loss margin.

The script was containerized in a Docker environment and deployed on the institution’s Kubernetes cluster. Each job was allocated one NVIDIA A100 GPU with 40 GB of graphics memory. The source code is made publicly available on GitHub: <https://github.com/MXGHarryLiu/HemoSight>.

##### Supervised fine-tuning

To assign classification labels, we constructed a support set from the training data by pairing embeddings generated by the pretext-trained encoder with their corresponding hematopathologists’ annotations. A linear support vector machine (SVM) classifier [7] was employed to map the embeddings to categorical labels. The SVM used a one-vs-rest decision function for multi-class classification, with input standardization and probability estimates enabled for output predictions. For subsequent analyses, the embedding space learned from the pretext training remained fixed while the SVM was retrained for specific tasks. To assess the impact of labeled data proportions on performance, we incrementally sampled labeled data stratified by cell type for inclusion in the support set. To account for class imbalance, we report macro-averaged scores henceforth.

##### Comparison with supervised baseline models

To establish a baseline for comparison with our self-supervised model, we implemented a basic supervised classifier using the same EfficientNetV2-B0 encoder. This architecture closely mirrored that of the self-supervised model, with the primary difference being a single encoder head and the addition of a dense layer with Softmax activation following the dropout layer to produce class probabilities for 9-way classification.

Following standard transfer learning practices, we employed a two-stage training strategy using the Adam optimizer, a batch size of 32, and sparse categorical cross-entropy loss. In the first stage, we trained the model with a learning rate of 0.001, freezing all layers except the top layer, for 10 epochs. In the second stage, we unfroze all layers except the batch normalization layers and fine-tuned the model for an additional 20 epochs at a reduced learning rate of ${10}^{-5}$. For datasets with fewer than or equal to 450 images, we adjusted the training protocol to mitigate overfitting and ensure convergence: the second stage was omitted, and the model was trained for 30 epochs with only the top layer unfrozen.

##### Investigating the model’s explainability using RISE

We adopted RISE (Randomized Input Sampling for Explanation [8]) to generate saliency maps to visually inspect the model’s decision. First, $N=5000$ random binary masks were generated at the size of 8 × 8 pixels with uniform probability of 0 or 1 for each pixel. The masks were up-sampled to 224 × 224 using bilinear interpolation before being applied to input images, which were first cropped and color-normalized. This generated $N$ masked images for each input image, sharing the same random masks. The masked images were passed to the feature encoder and SVM classifier to obtain class probability. The saliency maps were computed using the sum of the random masks weighted by the output probabilities of the respective class.

#### Web application packaging and deployment

To facilitate easy access to the model, we developed a full-stack web application comprising four Docker containers. The frontend, built with ReactJS and styled using the Bootstrap framework, is hosted in one container. RESTful APIs handle communication between the frontend and the backend container, which is implemented using FastAPI (version 0.105) in Python.

To enhance server responsiveness, a dedicated worker container was introduced to manage model inference. This container, also built with FastAPI, runs TensorFlow models in Python and interacts with a database container (MongoDB). It subscribes to a change stream from the database for incoming image classification requests and writes the processed results back to the database. This modular design allows for scalability, enabling multiple instances of the worker container to be deployed simultaneously to accelerate model inference.

The application visualizes results directly in the web browser as interactive Vega charts [9], providing an intuitive and responsive user experience.

### References

[1]	Acevedo A, Merino A, Alférez S, Molina Á, Boldú L, Rodellar J. A dataset of microscopic peripheral blood cell images for development of automatic recognition systems. Data Brief 2020;30:105474. https://doi.org/10.1016/J.DIB.2020.105474.

[2]	Tan M, Le Q V. EfficientNetV2: Smaller Models and Faster Training. Proc Mach Learn Res 2021;139:10096–106.

[3]	Deng J, Dong W, Socher R, Li L-J, Kai Li, Li Fei-Fei. ImageNet: A large-scale hierarchical image database 2010:248–55. https://doi.org/10.1109/CVPR.2009.5206848.

[4]	Ioffe S, Szegedy C. Batch Normalization: Accelerating Deep Network Training by Reducing Internal Covariate Shift. 32nd International Conference on Machine Learning, ICML 2015 2015;1:448–56.

[5]	Buchsbaum G. A spatial processor model for object colour perception. J Franklin Inst 1980;310:1–26. https://doi.org/10.1016/0016-0032(80)90058-7.

[6]	Schroff F, Kalenichenko D, Philbin J. FaceNet: A Unified Embedding for Face Recognition and Clustering. Proceedings of the IEEE Computer Society Conference on Computer Vision and Pattern Recognition 2015;07-12-June-2015:815–23. https://doi.org/10.1109/cvpr.2015.7298682.

[7]	Cortes C, Vapnik V, Saitta L. Support-vector networks. Machine Learning 1995 20:3 1995;20:273–97. https://doi.org/10.1007/BF00994018.

[8]	Petsiuk V, Das A, Saenko K. RISE: Randomized Input Sampling for Explanation of Black-box Models. British Machine Vision Conference 2018, BMVC 2018 2018.

[9]	Satyanarayan A, Moritz D, Wongsuphasawat K, Heer J. Vega-Lite: A Grammar of Interactive Graphics. IEEE Trans Vis Comput Graph 2017;23:341–50. https://doi.org/10.1109/TVCG.2016.2599030.
