## Supplementary figures and images for "Interpretable morphology mapping of peripheral blood leukocytes using annotation-efficient artificial intelligence"

### Supplementary Figure 1

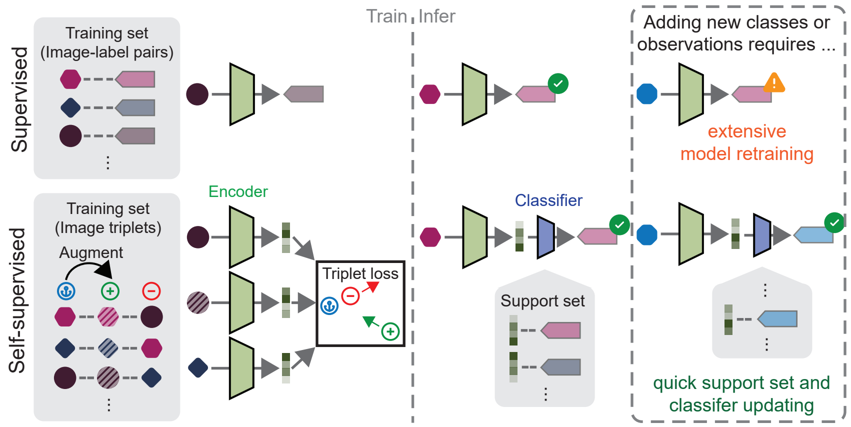

### Supplementary Figure 2

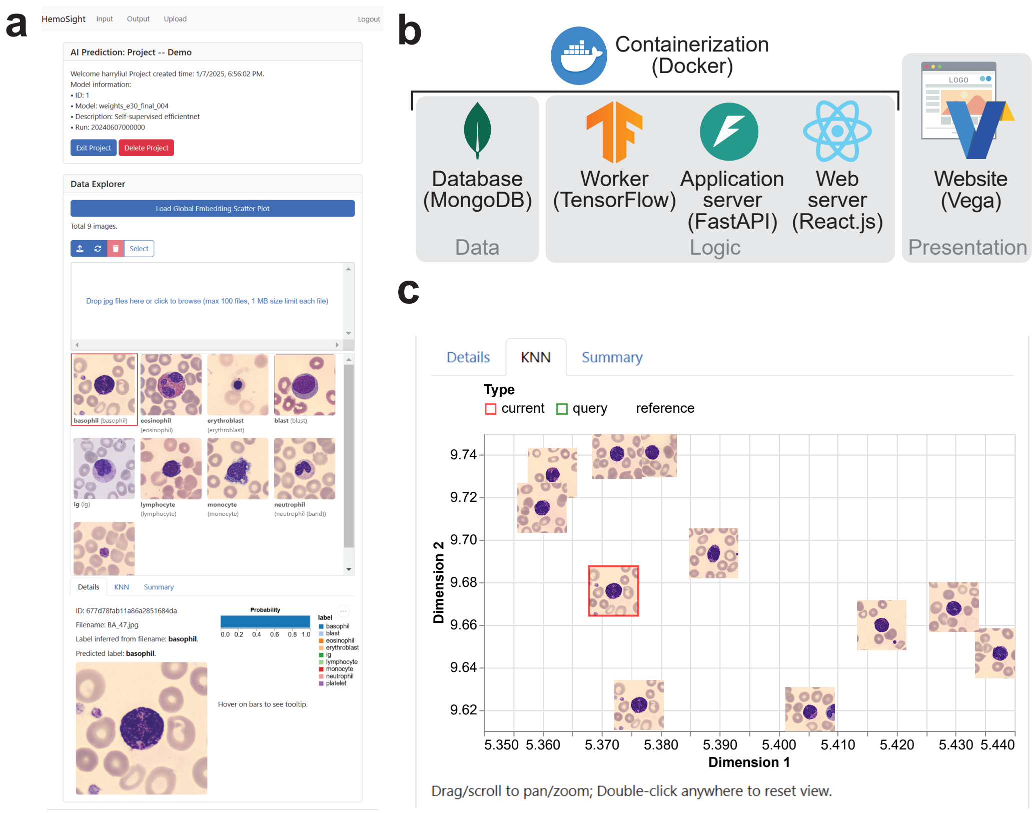
